# Cold sensing by a glutamate receptor drives avoidance behavior in *Drosophila* larvae

**DOI:** 10.1101/2025.10.05.680544

**Authors:** L. Amanda Xu, Hyun-Joon Son, Xu Bai, Ruonan Li, Elizabeth A. Ronan, Gun-Ho Kim, Bing Ye

**Author notes:** Corresponding Authors (B.Y.); (G.K.).

## Abstract

The ability to sense and avoid noxious environments is essential for animal survival; yet, how this is achieved at the behavioral, neuronal, and molecular levels is not well understood. Here, we use *Drosophila* larvae as a model to investigate how animals sense and avoid cold temperatures. By implementing custom-built thermoelectric devices capable of delivering rapid and precise thermal stimuli, we find that cold delivered to the larval head evokes robust escape behavioral responses. We identify a group of head-located cold-sensitive neurons as necessary and sufficient for such avoidance responses. We further demonstrate that the kainate-type glutamate receptor *Clumsy* acts as a novel cold sensor required for head cold sensitivity. Knockdown of *Clumsy* in head cold-sensing neurons suppresses their cold sensitivity. Heterologous expression of *Clumsy* confers cold sensitivity. Our results show that *Drosophila* larvae have evolved the capacity to detect and avoid cold temperatures through a previously uncharacterized cold-sensing mechanism.

## INTRODUCTION

The detection and avoidance of harsh environments is crucial to physiological homeostasis and survival. Noxious cues, such as noxious temperatures, pungent chemicals, and harsh mechanical forces, can trigger protective responses.^1^ Animals have evolved sophisticated sensory systems to detect such noxious stimuli that drive avoidance responses; yet, the underlying behavioral, neuronal, and molecular mechanisms are not well understood and remain a subject under intensive investigation. Temperature is a universal environmental cue affecting animals’ well-being. The ability to sense and avoid extreme temperatures is fundamental for survival.^1^ Temperature cues are first detected by molecular thermosensors expressed in temperature-sensitive neurons, which sense thermal stimuli and relay this information to the brain. While the mechanisms underlying noxious heat, innocuous heat, and innocuous cool sensation are well characterized, those underlying noxious cold sensation are less understood.

The fruit fly *Drosophila melanogaster* provides a powerful genetic system to dissect the molecular, neuronal, and behavioral mechanisms of thermosensation. Both the larvae and adults of *Drosophila* exhibit complex thermosensitive behaviors. One of the best-studied behaviors is thermotaxis, where flies seek favorable temperature zones (18 - 25°C) when placed on a thermal gradient.^2,3^ Thermosensitive behaviors are mediated by thermosensory neurons that detect innocuous warmth and cool temperatures, allowing flies to navigate towards their preferred temperature environments.^4^ In adults, warmth detection involves the Anterior Cells in the head that express dTRPA1, and Warm Cells in the arista expressing the ionotropic gustatory receptor Gr28b(D).^3,5,6^ Cool detection is mediated by aristal Cool Cells expressing Ionotropic Receptors IR21a, IR25a, and IR93a, as well as the Sacculus Cells, which may employ TRPP family members such as *Brivido1–3* or IRs in thermosensitivity.^7,8^ In larvae, Dorsal Organ Cool Cells (DOCCs), a trio of cool-sensing cells located in the dorsal organ ganglion (DOG), are required for cool detection.^9,10^ These neurons express the Ionotropic Receptors (IRs) IR21a and IR25a, which contribute to cool sensing.^9^ In addition to thermotaxis behaviors to innocuous temperatures, *Drosophila* also detect and respond to noxious heat, which is sensed by dendritic arborization (DA) neurons along the body wall that express the TRPA channel *painless*, essential for rolling responses, a type of avoidance behavior.^11–14^

By contrast, it remains unclear whether and how *Drosophila* avoid cold environments. Previous work found that global cooling of the entire larval body to cold temperatures evoked full-body contraction. In addition, simultaneous delivery of cold and touch stimuli to the larval body with a probe elicited full-body contraction and head and/or tail raise. As the larvae stayed immobile, these observations do not reveal a behavioral mechanism that allows the animals to escape the cold stimuli delivered. As such, it remains an open question whether *Drosophila* larvae have evolved the capacity to escape cold environments to promote their survival.

Here, we engineered customized thermoelectric devices capable of delivering thermal stimuli to localized body regions of *Drosophila* larvae without physical contact, enabling us to comprehensively interrogate whether, and how, larvae respond to cold temperatures. We found that cold stimuli delivered to the larval body evoke robust escape behavioral responses, with the head exhibiting the strongest response, enabling the animal to avoid cold environments. We discovered a new set of cold-sensitive neurons in the larval head that are essential and sufficient for head-specific cold avoidance behavior. Moreover, we identified the kainate-type glutamate receptor *Clumsy* as a novel type of cold sensor. Heterologous expression of *Clumsy* in mammalian cell lines conferred cold sensitivity, suggesting it is sufficient to function as a cold receptor. Our results demonstrate that cold evokes robust, ethologically relevant escape responses in *Drosophila* larvae, and uncover the underpinning neural and molecular mechanisms.

## RESULTS

### A new approach to studying acute, cold-responsive behaviors in *Drosophila* larvae

We first aimed to characterize whether and how *Drosophila* larvae respond to acute cold stimuli at the behavioral level. *Drosophila* larvae normally locomote forward via peristaltic body waves. We sought to deliver thermal stimuli to specific body areas of mid-third instar larvae during forward peristalsis: the head, mid-section, and tail. To apply localized cold stimuli, previous studies placed the tip of a pre-chilled probe in direct contact with the larval cuticle.^15^ However, this method lacks specificity because it introduces confounding mechanical input, making it difficult to distinguish cold-specific responses from mechanosensory responses evoked by physical contact (also see below). This limitation makes it especially difficult to assess thermosensory responses in the larval head, which is highly sensitive to touch and prone to exhibiting strong mechanosensory responses.

To overcome these technical challenges, we engineered a touchless thermoelectric device that can deliver localized thermal stimuli rapidly and precisely in the absence of mechanosensory input (Figures 1A and Figure S1). The probe tip was cooled to various set temperatures and positioned as close as possible, for efficient convectional and radiational thermal transfer without touching, three different regions of the larval body during locomotion—the head, mid-section, and tail (Figure 1A). *Drosophila* larvae thrive at around 25°C and temperatures near 18°C fall within the innocuous cool range, whereas temperatures below 15°C are generally considered noxious cold.^16^ The device was calibrated accordingly to cool the surrounding air near the larvae to cool (e.g., 18°C) and cold (e.g., 13°C) temperatures without physical contact, allowing us to identify cold-evoked, body region-specific behaviors in aversive thermal environments (see methods).

**Figure 1.**
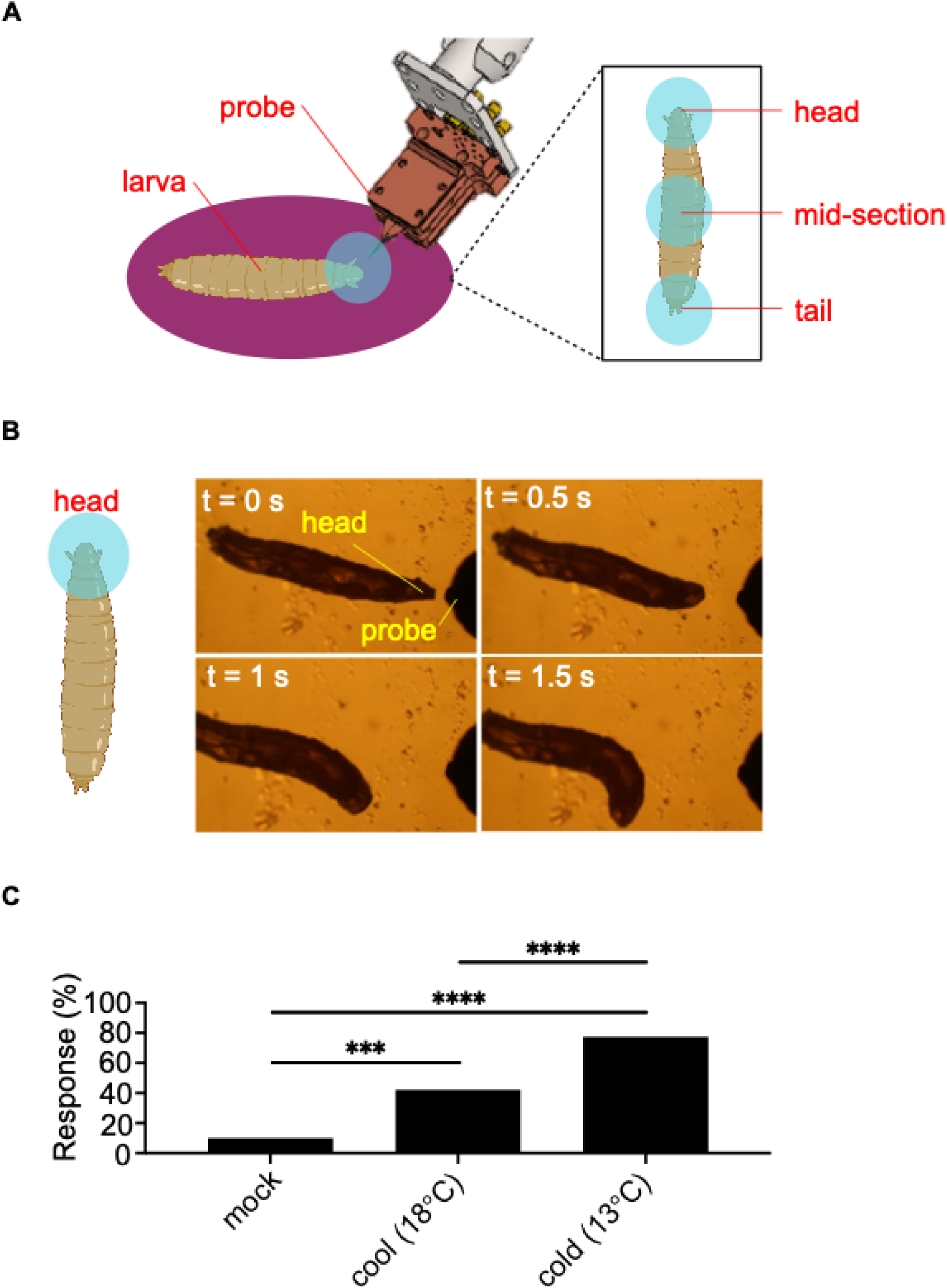
The head of *Drosophila* larvae responds robustly to cold. **(A)** Schematic showing the temperature-based behavioral assay. The pre-chilled probe tip was directed to the head, abdominal mid-section, and tail regions of the larval body to cool the surrounding air without physical contact. **(B)** Representative images of the head avoidance behavioral sequence. **(C)** Mock (ambient temperature); cool (18°C); and cold (13°C). The frequency of head avoidance responses increases significantly at colder temperatures. Chi-squared test. Wild-type Canton-S larvae were tested at room temperature control (n = 100), cool (n = 109), and cold (n = 71) conditions. ***p<0.001 and ****p<0.0001.

### The head of *Drosophila* larvae responds robustly to low temperatures

We found that the head of the larva exhibited the most robust responses to cold. Upon exposure to low temperature environments, the larva ceased forward locomotion. Then, the head withdrew and turned away from the probe, resulting in a directional change that allowed the larva to avoid the thermal stimuli and navigate to a new environment. We termed this behavior head avoidance (Figure 1B and Video S1). Cool stimuli (18°C) induced a modest head avoidance response, whereas colder temperatures (13°C) elicited much stronger responses, indicating heightened head sensitivity at lower temperatures (Figure 1C).

When directing the probe toward the larval tail during peristalsis, no notable cold-evoked behavioral responses were observed (Figure S2A and Video S2). Similarly, responses from the larval body wall, though detectable, were very weak (Figure S2B). Specifically, when targeting the mid-abdominal segments, a small proportion of larvae (∼5%) exhibited evasion behaviors during ongoing peristaltic movement at 13°C, with a modest increase to ∼25% at 9°C (Figure S2C). We defined this response as mid-section avoidance (Figure S2A and Video S2). Due to technical limitations, we were unable to assess cold-specific responses to temperatures below 9°C without inducing physical contact. This is because, although the surface of the probe tip could be set to subzero temperatures to deliver air temperatures lower than 9°C to the larva, such a setting causes substantial amounts of ice to accumulate on the tip of the probe, limiting its cooling capacity. Interestingly, when the surface of the probe tip was cooled to 9°C and physically contacted the larval mid-section, the mid-section avoidance response to 9°C was much more robust than that elicited by a non-contacting cold probe (Figure S2C). As contact with an ambient-temperature probe did not elicit a noticeable mid-section avoidance response (Figure S2C), this indicates that the cold and mechanosensory inputs may act synergistically to amplify avoidance responses. This underscores the importance of delivering pure thermal stimuli for accurate assessments of thermosensory behaviors.

These findings suggest that *Drosophila* larvae are most sensitive to cold temperature applied to their head. Therefore, we focused the remainder of our investigation on head-specific cold sensation.

### Neurons required for head cold sensation in *Drosophila* larvae

We then set out to identify the sensory neurons necessary for cold-elicited head avoidance. We first examined the Dorsal Organ Cool Cells (DOCCs), a trio of cool-sensing neurons expressed in the dorsal organ ganglion (DOG) on each side of the larval head, previously shown to mediate larval temperature preference in thermotaxis.^9,10^ To do so, we tested the head responses of larvae to cold while optogenetically inhibiting DOCCs via *Guillardia theta* anion channelrhodopsin-1 (GtACR1).^17,18^ GtACR1 was selectively expressed in DOCCs using a DOCCs-specific GAL4 driver, *R11F02*-GAL4. Transgenic larvae were then illuminated with 527-nm green light to activate GtACR1, and their head avoidance behavior in response to the cold probe was recorded and quantified. DOCCs-inhibited larvae responded normally to low temperatures, suggesting that they are not required for head sensation of acute cold (Figure 2A).

**Figure 2.**
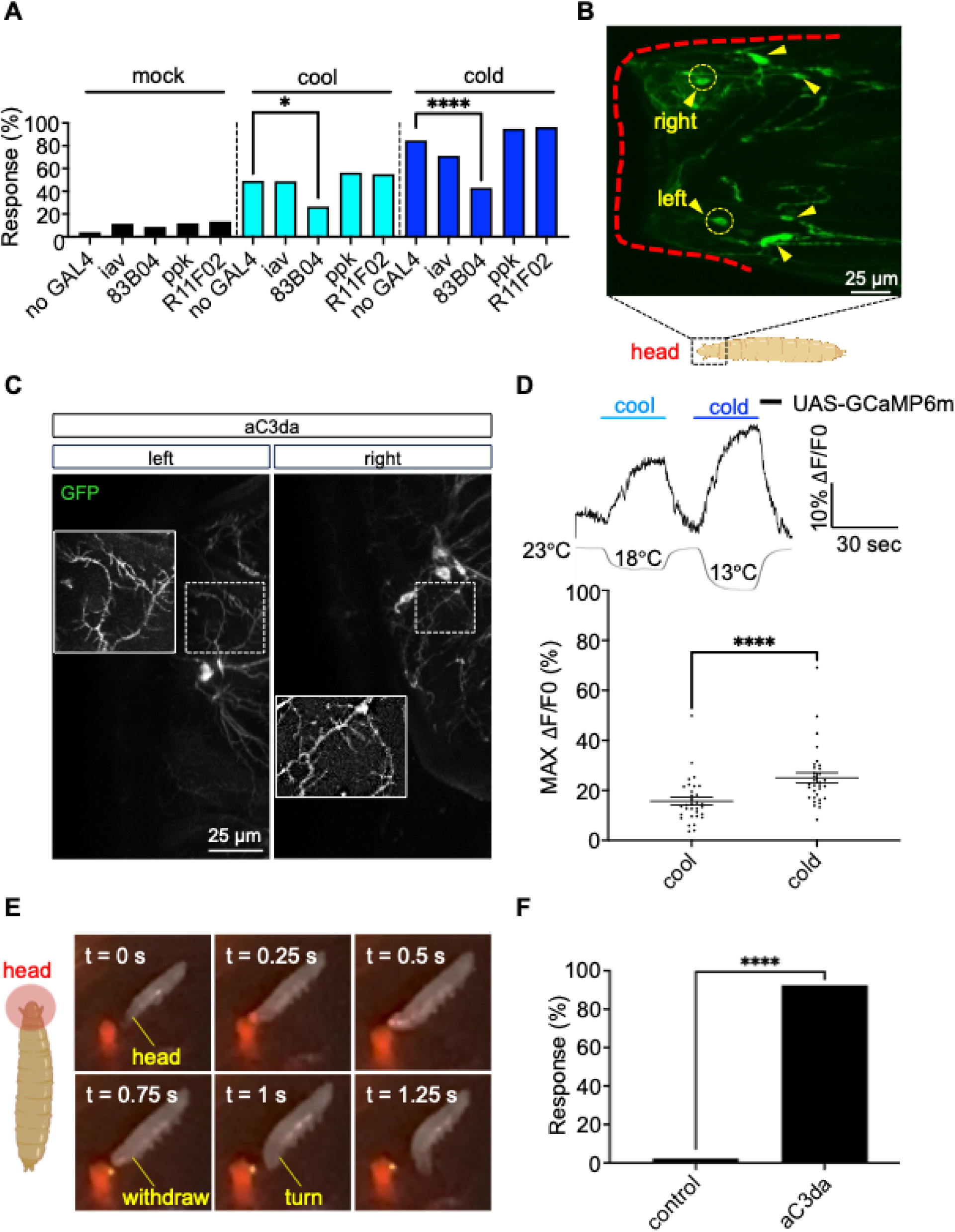
A group of neurons are important for cold sensitivity in the head of *Drosophila* larvae. **(A)** Optogenetic inhibition of 83B04-GAL4 labeled neurons reduces the frequency of head avoidance behavior at cool and cold temperatures. Since GtACR1 was expressed in each candidate neuron group via a cell-type-specific GAL4 driver, larvae expressing GtACR1 only without a GAL4 driver served as the negative control. *iav*-GAL4: Cho-specific driver; 83B04-GAL4: C3da-specific driver; *ppk*-GAL4: C4da-specific driver; *R11F02*-GAL4: DOCCs-specific driver. Chi-squared test. Bar graph, n ≥ 40 per genotype. *p<0.05 and ****p<0.0001. **(B)** Representative UAS-mCD8-GFP-mediated visualization of head 83B04-GAL4-positive neurons in an immobilized mid-third-instar larva. **(C)** Immunostaining of GFP-expressing aC3da (anterior C3da) neurons under 83B04-GAL4. Dendritic spikes, which are characteristic of C3da neurons, are magnified. Left: left anterior C3da cluster. Right: right anterior C3da cluster. **(D)** Top: Representative trace of cold and cool-induced calcium transients in GCaMP-expressing aC3da neurons. Length of the temperature label above the trace corresponds to the duration of cool/cold stimulation. Bottom: Maximum ΔF/F_0_ of individual aC3da neuronal clusters is plotted on the graph and used for statistical analyses (each dot indicates one cluster). N = 32 at both temperatures tested. Paired Student’s t-test. Error bars represent SEM. ****p<0.0001. **(E) and (F)** Optogenetic activation of 83B04-GAL4 labeled (aC3da) neurons in the larval head via head-localized illumination elicits rapid head avoidance behavior. Since ATR is necessary for optogenetic manipulation, larvae grown on food without ATR and subjected to the same illumination conditions served as the negative control (F). Chi-squared test. Bar graph, n = 43 (control) and n = 40 (anterior C3da activation). ****p<0.0001.

We then assessed chordotonal (Cho) neurons, which are low-threshold mechanosensory neurons implicated in proprioception and hearing.^19^ Cho neurons are expressed in the larval head next to the dorsal organ, in addition to their innervation along the body wall.^20^ We found that inhibition of neurons labeled by the Cho-specific driver *iav*-GAL4 did not affect avoidance responses in the head (Figure 2A).

Next, we tested class III and class IV dendritic arborization neurons (i.e., C3da and C4da neurons). Class III dendritic arborization (C3da) neurons were initially found to sense innocuous mechanical stimuli along the body wall,^21^ but were later reported to also be involved in cold sensing.^15^ Class IV dendritic arborization (C4da) neurons are polymodal nociceptors along the body wall mediating larval escape responses to noxious heat and nociceptive stimuli.^11,12^ C4da and C3da are well-characterized body wall neurons, with dendritic arbors across the body wall and axon terminals projecting into the ventral nerve cord (VNC). Yet, this does not exclude the possibility that their respective GAL4 drivers also label neurons in the head, or their potential involvement in head sensory processing.

We found that while inhibition of C4da neurons (labeled by *ppk*-GAL4) did not influence head avoidance responses (Figure 2A), inhibition of neurons labeled by the C3da-specific driver 83B04-GAL4 significantly decreased both cool- and cold-elicited head avoidance responses from those of no GAL4 controls (Figure 2A). These results demonstrate that these 83B04-positive neurons are essential for head cold sensation.

83B04-GAL4 labels C3da neurons located along the *Drosophila* body wall, yet the requirement of 83B04-GAL4-positive neurons for head avoidance to cold suggests that the 83B04-GAL4 driver may be labeling uncharacterized neurons in the head. To assess for the presence of head-localized neurons, we expressed membrane-bound GFP (UAS-mCD8-GFP) driven by 83B04-GAL4 and examined GFP-positive cells in the larval head. We found four bilaterally symmetrical neuronal clusters in the head region, including a distinct bilateral cluster located closest to the tip of the larval head, positioned to be among the first neurons activated by head-directed stimuli (Figure 2B). To further assess the morphology of the tip-most neuronal cluster, we performed immunostaining with an anti-GFP antibody in larvae expressing the UAS-CD4-tdGFP^vk33^ transgene under 83B04-GAL4 (Figure 2C). The dendritic arbors of the neurons displayed morphology characteristic of body wall C3da neurons, including short terminal branchlets projecting from main dendrites with minimal overlap (Figure 2C).^22^ Based on location and morphology, we classified these neurons as anterior C3da (aC3da).

### aC3da neurons are cold-sensitive

The requirement of aC3da for cold-evoked head avoidance and the fact that they are located on the cuticle surface suggest that they may be sensory neurons for cold. To test this, we developed a dissection protocol to target head sensory neurons for imaging (Figure S3A and see methods). We expressed the genetically encoded calcium indicator GCaMP6m under control of 83B04-GAL4 and performed calcium imaging in larval head preparations in which only aC3da neurons were present. We then monitored calcium transients in aC3da neurons in response to sequential cool and cold stimuli delivered using a thermoelectric pad device (Figures S1B and S3B). We found that cooling evoked robust calcium activity in aC3da neurons, with cold temperature (13°C) eliciting a stronger response than cool temperature (18°C) (Figure 2D). These results demonstrate that the aC3da neurons are indeed cold-sensitive.

### aC3da neurons are sufficient for head avoidance behavior

Having established the cold-sensitivity of aC3da neurons and their necessity for cold-induced head avoidance behavior, we wondered if these neurons are also sufficient to elicit head avoidance responses in larvae. To test this possibility, we optogenetically stimulated C3da neurons by expressing CsChrimson driven by 83B04-GAL4. To achieve head-specificity in C3da activation, we selectively illuminated the head of transgenic larvae with a small-diameter 655-nm red light spot to activate CsChrimson in head-localized C3da, and recorded their behavior in response to optogenetic activation. Indeed, we found that selectively activating head C3da was sufficient to elicit robust head avoidance responses, where forward locomoting larvae withdraw the tip of their head, followed by a turn leading to directional change (Figures 2E and 2F; Video S4). Interestingly, selective activation of C3da neurons in other larval body regions elicited unique behavioral responses distinct from those exhibited by the head (Figure S4 and Videos S5-S7). Selectively illuminating the larval abdominal mid-section during peristalsis induced mid-section avoidance behavior similar to responses exhibited to cold (Figure S4A and Video S5), whereas selective illumination of the tail evoked no observable responses (Figure S4B and Video S6). In contrast, global activation of C3da via illumination of the entire larval body elicited full-body contraction behavior, where both the head and tail withdraw simultaneously (Figure S4C and Video S7). Our results indicate that the activation of C3da neurons in different larval body regions produces distinct behavioral responses, but only selective stimulation of aC3da neurons in the larval head elicits head avoidance behavior.

### The kainate-type glutamate receptor Clumsy mediates head cold-avoidance behavior

To identify the cold-sensing receptors that mediate head cold sensitivity, we screened candidate thermoreceptor-encoding genes for defects in head cold avoidance. We first examined ionotropic receptor IR25a, a cool sensor expressed in DOCCs necessary for larval thermotaxis.^9^ *Ir25a*^1^ mutants and IR25a knockdown in 83B04-positive neurons did not result in behavioral defects to cold, suggesting IR25a does not play an important role in aC3da-mediated cold-sensing (Figures 3A and S5A). We then assessed the TRP channels NompC, Trpm, and Pkd2, which are previously implicated as noxious cold sensors in body-wall C3da neurons.^15^ Knockdown of these channels in C3da neurons by RNAi did not affect head avoidance responses, suggesting that they are not essential for cold sensation in the larval head (Figure S5A).

**Figure 3.**
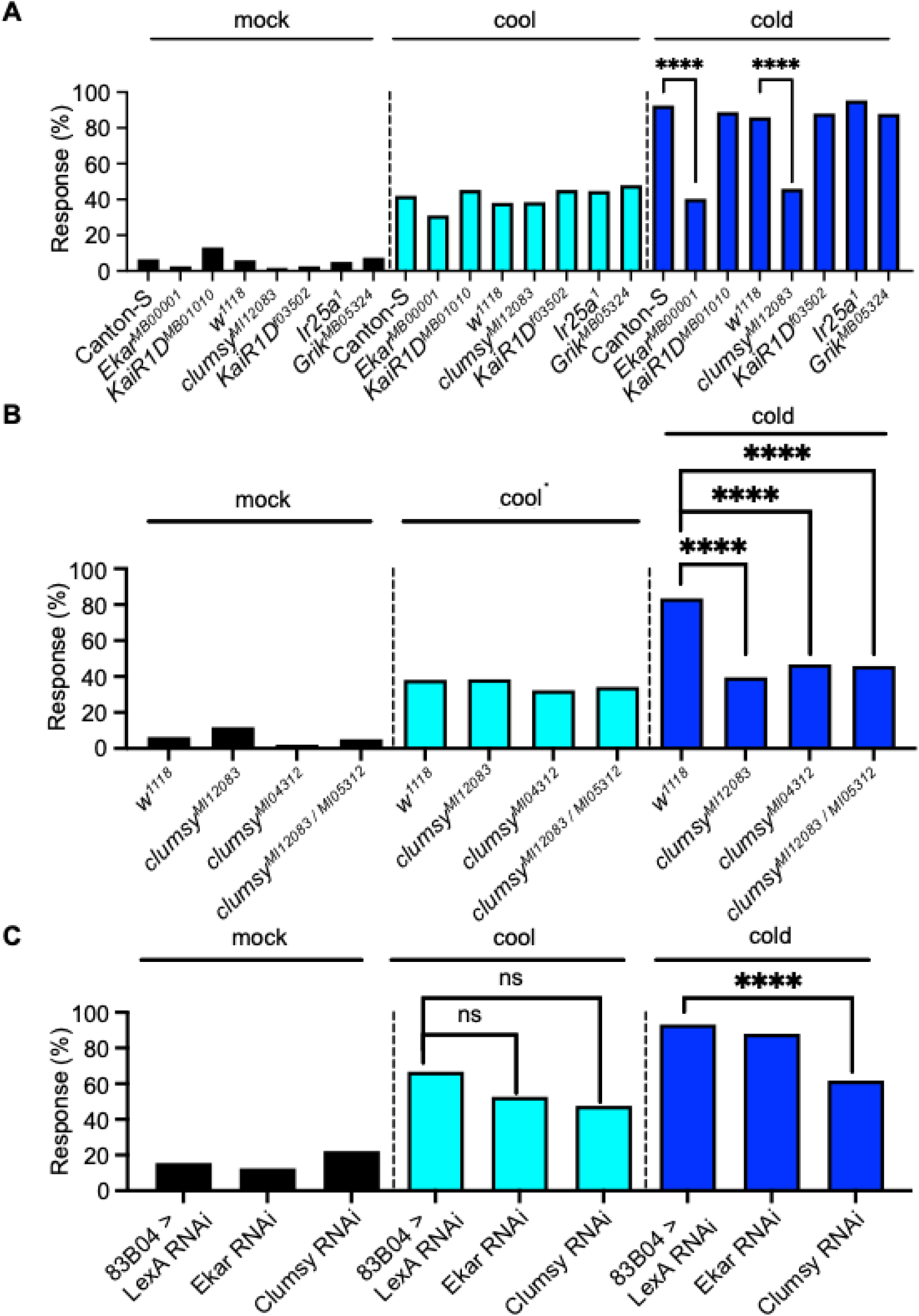
The kainate-type glutamate receptor *Clumsy* mediates head cold avoidance behavior in aC3da neurons. **(A)** Head avoidance responses of genetic mutants to cool and cold temperatures. *Ekar* and *Clumsy* genetic mutants are defective in sensing cold, but not cool, temperature. Each genetic mutant was compared to its corresponding genotypic background (either Canton-S or *w*^118^). Chi-squared test. Bar graph, n > 40 per genotype. ****p<0.0001. **(B)** The head of *Clumsy* homozygous and transheterozygous mutants are defective in sensing cold, but not cool, temperature. Mutant responses were compared with those of *w*^118^, its genotypic background. Chi-squared test. Bar graph, n > 40 per genotype. ****p<0.0001. **(C)** RNAi-mediated knockdown of *Clumsy*, but not *Ekar*, in aC3da neurons leads to a head cold-sensing phenotype. A shRNA against LexA (LexA RNAi) was used as the negative control. Chi-squared test. Bar graph, n > 40 per genotype. ****p<0.0001.

We then tested kainate-type glutamate receptors, a new type of cold sensor recently discovered in *C. elegans* and mice.^23,24^ The *Drosophila* genome encodes five neuronal kainate receptors: KaiR1D, Grik (KaiR1C), Ekar, Clumsy, and CG11155.^25^ We examined mutants of *KaiR1D*, *Grik*, *Ekar*, and *Clumsy* (mutants of *CG11155* are not available), and C3da-specific RNAi knockdowns of *KaiR1D* and *CG11155*. Mutants or knockdowns of *KaiR1D*, *Grik*, and *CG11155* showed no behavioral phenotypes (Figures 3A and S5A). In contrast, *Ekar^MB^*^00001^ and *Clumsy^MI^*^12083^ homozygous mutants displayed a significant defect in cold-evoked head avoidance responses to cold, but not cool temperature (Figure 3A). A similar cold-specific phenotype was observed in another allele, *Clumsy^MI^*^05312^, and in *Clumsy^MI^*^12083^/*Clumsy^MI^*^05312^ transheterozygous mutants (Figure 3B). The head of *Clumsy* mutants responds normally to heat and touch stimuli (Figure S5B), confirming that their head avoidance phenotype is specific to cold sensation.

In contrast, although the head avoidance phenotype to cold persisted in other homozygous and transheterozygous mutants of *Ekar* (Figure S5C), the mutants were also defective in sensing heat and touch (Figure S5D). The non-specific head avoidance phenotype suggests that the *Ekar* mutation may be affecting downstream neural processing in the head rather than the initial detection of cold stimuli, leading to a common behavioral defect in response to different types of head-directed stimuli. Therefore, our subsequent investigations focused on the cold-specific phenotype of *Clumsy*.

To determine whether *Clumsy* is required in aC3da neurons to mediate cold-sensing, we selectively knocked down *Clumsy* in aC3da by expressing UAS-*clumsy* RNAi with 83B04-GAL4. Strikingly, as was the case with the mutant larvae, *Clumsy* knockdown in the aC3da led to a defect in sensing cold, but not cool, temperatures in the head avoidance assay (Figure 3C). In contrast, RNAi-mediated knockdown of *Ekar* in the same neurons did not reproduce the cold-avoidance defect (Figure 3C). These results suggest that *Clumsy* functions in the aC3da neurons to mediate cold sensing behavior in *Drosophila* larvae.

### Clumsy is required for the cold sensitivity of aC3da

We next determined whether Clumsy is required for aC3da to detect cold by using calcium imaging. We first assessed how *Clumsy* knockdown in aC3da neurons, as well as whole-body *Clumsy* mutations, affected their thermosensitivity to sequential cool and cold stimuli in larval head preparations (Figures S3A and S3B; see methods).

aC3da neurons exhibited robust calcium responses to both cool and cold, indicating that they are thermosensitive across a range of lower temperatures (Figures 2E and 2F). Strikingly, two independent *Clumsy* mutant alleles showed reduced calcium responses to cold (Figures 4A and 4B). Similarly, RNAi-mediated knockdown of *Clumsy* in aC3da neurons led to a significant reduction in cold-evoked calcium transients (Figures 4C and 4D). In contrast, cool-evoked responses were largely unaffected. Only one of the two *Clumsy* mutant alleles showed a modest reduction in response to 18°C (Figure 4B), while RNAi knockdown did not significantly alter cool sensitivity (Figure 4D). These results suggest that *Clumsy* is specifically required for aC3da neuronal responses to cold, but not cool, temperatures.

**Figure 4.**
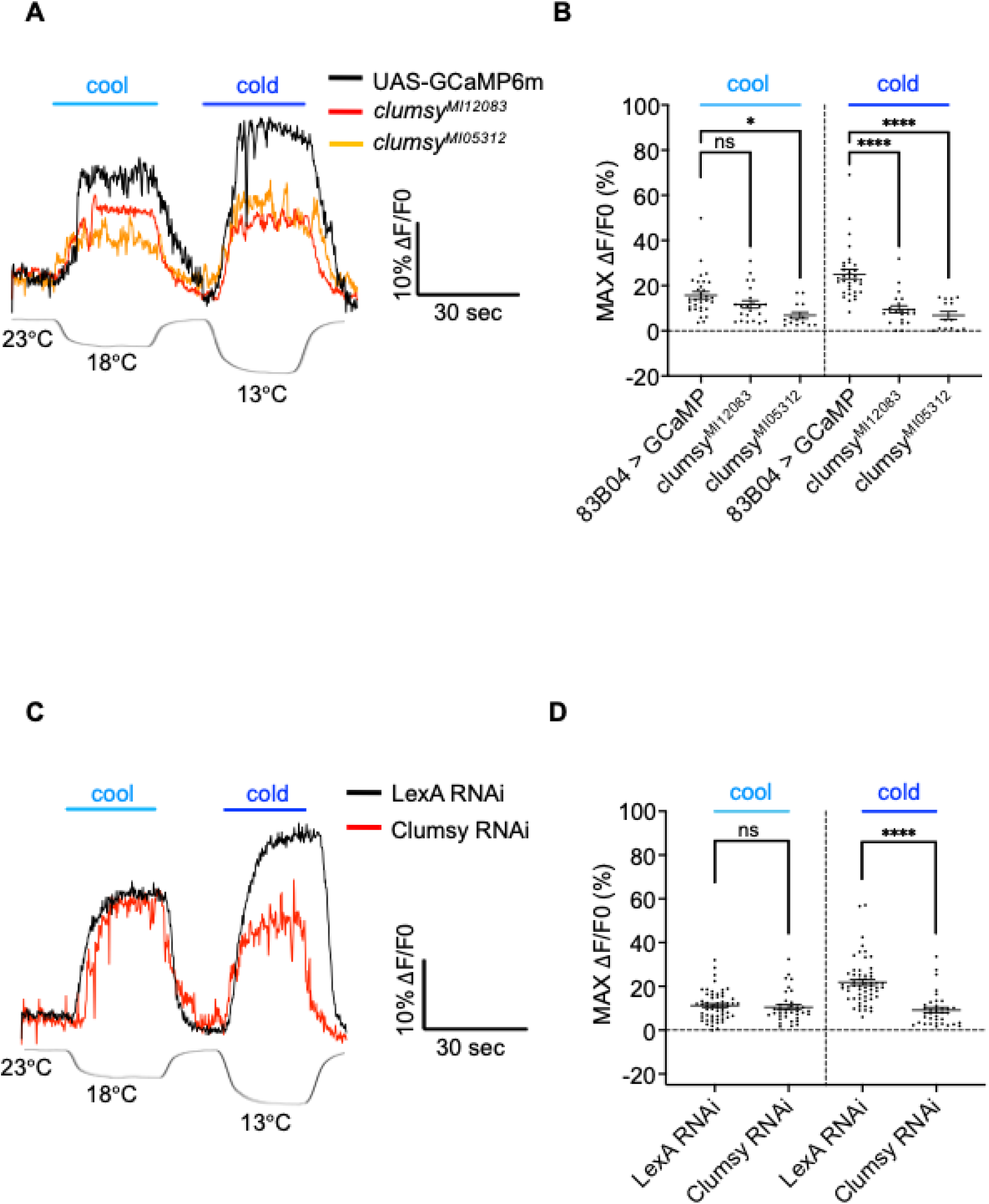
aC3da neurons require Clumsy to sense cold. **(A)** Representative traces of cool and cold-induced calcium transients of aC3da neurons in GCaMP-expressing controls and *Clumsy* mutants. Length of the temperature label above the trace corresponds to the duration of cool/cold stimulation. **(B)** Maximum ΔF/F_0_ of individual aC3da neuronal clusters is plotted on the graph and used for statistical analyses (each dot indicates one cluster). At both tested temperatures, control (n = 32), *Clumsy^MI^*^12083^ (n = 22), *Clumsy^MI^*^05312^ (n = 14). One-Way ANOVA with Tukey multiple comparisons post hoc analysis. Error bars represent SEM. *p<0.05 and ****p<0.0001. **(C)** Representative traces of cool and cold-induced calcium transients of aC3da neurons in GCaMP-expressing controls and with *Clumsy* RNAi-mediated knockdown. Length of the temperature label above the trace corresponds to the duration of cool/cold stimulation. **(D)** Maximum ΔF/F_0_ of individual aC3da neuronal clusters is plotted on the graph and used for statistical analyses (each dot indicates one cluster). N = 56 for control and n = 36 for clumsy RNAi at both tested temperatures. Unpaired Student’s t-test. Error bars represent SEM. ****p<0.0001.

Since larvae in the imaging assay were subjected to sequential thermal stimuli—first at 18°C and then at 13°C (Figure S3B)—we considered the possibility that reduced responses to cold in *Clumsy* mutants and RNAi knockdowns might reflect sensory desensitization due to prior cool exposure. To assess this possibility, we repeated the experiment using a modified stimulation protocol in which a single cold stimulus (13°C) was applied without prior exposure to cool (18°C) (Figure S3C). Notably, the cold-sensing deficits in both *Clumsy* mutants (Figures S6A and S6B) and RNAi knockdowns (Figures S6C and S6D) persisted under this single-stimulus protocol, confirming that the cold-sensing phenotype is due to *Clumsy* function and not desensitization. Together, these data demonstrate that aC3da neurons are sensitive to both cool and cold temperatures, and that *Clumsy* is specifically required for their cold—but not cool—responses. This finding further supports a direct role for *Clumsy* in cold sensing at the neuronal level.

### Clumsy functions as a cold-sensing receptor

The requirement of *Clumsy* for the cold responses of aC3da raised the question of whether *Clumsy* can directly function as a cold receptor. One of the most common methods to assess this is to express the candidate receptor-encoding gene in cold-insensitive heterologous cells and examine whether cold sensitivity could be conferred to these conventionally non-cold-sensing cells. To test whether *Clumsy* can function as a cold sensor, we expressed the RA and RB splicing isoforms of *Clumsy* in Chinese Hamster Ovary (CHO) cells and performed Fura-2 ratiometric calcium imaging of CHO cells during exposure to cool (Figure S7A) and cold (Figure S7B) temperatures. While CHO cells are insensitive to cold, transfection of both *Clumsy* isoforms in CHO cells conferred robust sensitivity to cold temperature (Figures 5A and 5B). By contrast, we observed little to no cool sensitivity conferred to *Clumsy*-expressing CHO cells at 18°C (Figures 5C and 5D), consistent with the finding that the activation threshold of this type of cold sensor is around 18°C^23,24^ and primarily responds to cold temperatures.^23,24^ Together, these results demonstrate that *Clumsy* is sufficient to sense cold, strongly suggesting that *Clumsy* is a cold-sensing receptor.

**Figure 5.**
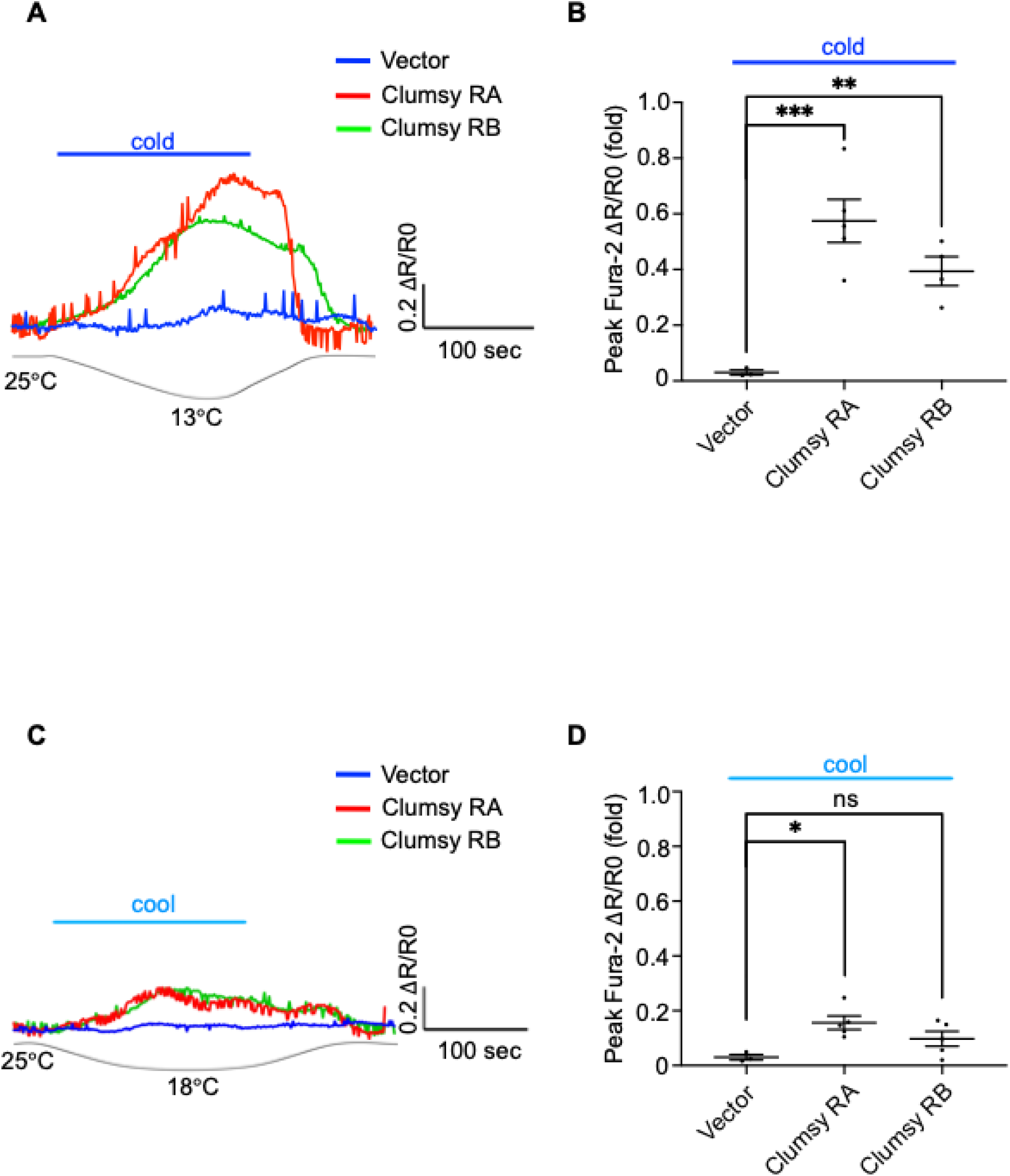
Clumsy confers cold sensitivity to CHO cells. **(A)** Representative traces of cold-induced calcium transients in CHO cells transfected with *Clumsy*, visualized via Fura-2 imaging. Length of the temperature label above the trace corresponds to the duration of cold stimulation. **(B)** Peak ΔR/R_0_ of cold-evoked calcium responses of *Clumsy*-expressing CHO cells are plotted on the graph and used for statistical analyses (each dot indicates one coverslip containing at least 20 CHO cells). Vector (n = 3 [at least 60 cells]), Clumsy RA (n = 5 [at least 100 cells]), Clumsy RB (n = 4 [at least 80 cells]), One-Way ANOVA with Dunnett’s multiple comparisons post hoc analysis. Error bars represent SEM. **p<0.01 and ***p<0.001. **(C)** Representative traces of cool-induced calcium transients in CHO cells transfected with *Clumsy*, visualized via Fura-2 imaging. Length of the temperature label above the trace corresponds to the duration of cool stimulation. **(D)** Peak ΔR/R_0_ of cool-evoked calcium responses of *Clumsy*-expressing CHO cells are plotted on the graph and used for statistical analyses (each dot indicates one coverslip containing at least 20 CHO cells). Vector (n = 3 [at least 60 cells]), Clumsy RA (n = 5 [at least 100 cells]), Clumsy RB (n = 5 [at least 100 cells]), One-Way ANOVA with Dunnett’s multiple comparisons post hoc analysis. Error bars represent SEM. *p<0.05.

## DISCUSSION

How animals detect and avoid unfavorable environments is a fundamental question in neuroscience. In this study, using *Drosophila* larvae as a model, we show that cold temperatures evoke a robust escape response, indicating that these animals are capable of sensing and avoiding cold environments. This behavioral response would be beneficial to these animals, as it provides a mechanism for them to avoid and survive unfavorable environments. We show that distinct clusters of neurons in the head, aC3da, are activated by cold stimuli, and required for cold-evoked head avoidance behavior. Optogenetic activation of these neurons is sufficient to recapitulate cold-evoked head avoidance responses. We further demonstrate that the kainate-type glutamate receptor *Clumsy* is cold-sensitive and involved in cold, but not cool, detection by aC3da neurons. Our findings offer new insights into the behavioral, neuronal and molecular basis of cold detection and avoidance in *Drosophila* larvae.

Our behavioral analysis uncovered pronounced differences in cold responsiveness along the anterior-posterior axis of the larval body. Head avoidance was the most robust and consistent response to cold, whereas mid-section and tail regions were less responsive. This difference in cold sensitivity suggests that distinct cold sensory mechanisms operate across different regions of the larval body. The heightened cold sensitivity of the head may reflect its greater ethological relevance: during navigation and foraging, *Drosophila* larvae primarily interact with their environment head-first, using the anterior end to explore their surroundings. Thus, cold-evoked behaviors initiated through the head are likely more ethologically relevant for the larvae to turn away from noxious cold to explore a new environment more favorable to their survival. Understanding the sensory and molecular mechanisms driving head-specific cold avoidance may thus offer more ethologically meaningful insights into how larvae detect and respond to environmental threats.

Our characterization of the aC3da neurons provides new insight into the somatosensory organization and behavioral adaptation of *Drosophila* larvae. The 83B04-GAL4 driver labels a previously uncharacterized group of head sensory neurons, which we examined both morphologically and functionally. We found that 83B04-positive neurons in the head exhibit the characteristic morphology of body-wall C3da neurons, and we thus classified them as anterior C3da (aC3da) neurons. Optogenetic inhibition of these neurons significantly attenuates cold-elicited head avoidance behavior, while *in vivo* imaging revealed robust cold-evoked calcium activity in aC3da, suggesting that these neurons are key mediators of head cold sensitivity. In addition, optogenetic activation of aC3da is sufficient to drive robust head-specific avoidance responses. Such avoidance behavior is likely more advantageous for larval survival than full-body contraction, as it enables the animal to actively escape from unfavorable environments rather than remain immobile. The sufficiency of aC3da for this adaptive response underscores the ecological relevance of aC3da function.

Interestingly, knockdown of the TRP channels *NompC*, *Trpm*, and *Pkd2*—despite their proposed role in cold nociception^15^—did not affect head cold sensitivity in our assay (Figure S5A). One explanation might be the different behavioral assays used in the present and previous studies. Previous studies placed the thermal probe on the larval dorsal midline, typically at the abdominal segment 4, for either 10 seconds or until the larva exhibits its first response. The mechanical component of the stimuli may confound the behavioral responses. Given that these TRP channels are also mechanosensors, their mutant behavioral phenotypes may reflect combined mechanosensory and thermosensory defects. Our study also suggests that IR25a, previously implicated in DOCC-mediated thermotaxis,^9^ does not contribute significantly to acute cold-avoidance behavior. Neither mutation nor RNAi knockdown of IR25a affected larval head responses to cold stimuli (Figures 3A and S5A). This unexpected result may also reflect differences in behavioral strategies: thermotaxis involves sustained locomotor decision-making over extended periods, whereas head cold avoidance represents an acute response to cold stimuli. Thus, distinct receptors may underlie rapid versus long-term cold detection in the *Drosophila* larval head sensory system.

*Clumsy* expression in heterologous cells is sufficient to confer robust sensitivity to cold, but not cool, temperature (Figure 5), a result consistent with the cold-specific function of *Clumsy*. *In vivo* calcium imaging showed that *Clumsy*-deficient aC3da respond normally to cool but less to cold (Figure 4), aligning with behavioral phenotypes observed in genetic mutants and RNAi knockdown of *Clumsy* (Figure 3). These results indicate that, while *Clumsy* is essential for cold sensing in aC3da, it is not required for the cool sensitivity of these neurons, suggesting that different mechanisms underlie cool and cold sensation. Furthermore, the presence of cool-induced responses and residual cold sensitivity in both RNAi knockdowns and mutants of *Clumsy* raises the possibility that other, yet unidentified, ion channels or receptors may contribute to cool detection in the *Drosophila* head.

Our findings align with a growing body of evidence that kainate-type glutamate receptors, chemoreceptors conventionally associated with excitatory neurotransmission in the CNS,^26,27^ may also function as cold-sensing receptors in the periphery.^23,24^ This concept parallels discoveries that gustatory and chemoreceptors, such as Gr28b(D) and IRs, have unexpected thermosensory functions.^6,9^ Whether this functional versatility reflects ancient evolutionary roles or more recent adaptations remains an open question. Nevertheless, our study provides an entry point to better understand the molecular and neural mechanisms underlying cold sensation, and how these sensory systems contribute to behavioral responses to aversive thermal environments.

## MATERIALS AND METHODS

### *Drosophila* mutants and transgenic lines

Canton-S, *w*^1118,28^ UAS-mCD8::GFP,^29^ UAS-GtACR1,^18^ *ppk-*GAL4,^30^ *iav*-GAL4, 83B04-GAL4,^31^ *R11F02*-GAL4,^10^ UAS-CD4-tdGFP^vk^^33^, UAS-LexA RNAi, UAS-KaiR1D RNAi, UAS-Pkd2 RNAi, UAS-CG11155 RNAi, UAS-Ir25a RNAi, UAS-NompC RNAi, UAS-Trpm-RNAi, UAS-Ekar RNAi, UAS-_Clumsy RNAi, *Ekar*_*MB00001*_, *Ekar*_*MI02500-GFP*_, *Ekar*_*MI02500*_, *Ekar*_*MI09564*_, *KaiR1D*_*MB01010*_, *KaiR1D*_*f03502*, *clumsy^MI^*^12083^, *clumsy^MI^*^05312^, *Grik^MB^*^05324^, *Ir25a*^1^, UAS-GCaMP6m

### Generation of DNA Constructs

We generated constructs for CHO cell transfection by amplifying the *Clumsy* RA and *Clumsy* RB coding sequences via PCR and subcloning them into the pcDNA3.1-IRES-mCherry vector using NheI and XhoI restriction sites to produce pcDNA3.1-RA-IRES-mCherry and pcDNA3.1-RB-IRES-mCherry. The digested PCR product and vector were ligated with T4 DNA ligase, transformed into E. coli DH5α, and verified by Sanger sequencing.

### Thermoelectric devices

#### Probe device

For localized temperature control, a nickel-plated tungsten probe (72TN-J3, American Probe & Technologies, USA) was employed as the temperature-regulating element. The probe was inserted 15 mm into a five-sided copper component coupled with two thermoelectric modules (1MDL06, TEC Microsystems, Germany), allowing effective heat exchange along its shaft. A 0.5-mm-diameter thermistor (GA10K3MCD1, TE Connectivity, USA) was positioned 5 mm from the probe tip to provide temperature feedback. To maintain cooling efficiency and ensure stable temperature sensing, the probe was insulated along its length except for the distal 2 mm near the tip. The thermistor and thermoelectric modules were connected to a temperature controller (Newport, Model 3700, USA), which regulated the probe temperature. Heat generated on the hot side of the thermoelectric modules was dissipated by two water blocks. A temperature-controlled circulator chiller (Fisher Scientific Isotemp 6200 R28, USA) maintained the coolant of deionized water at 10 °C to provide stable heat exchange during operation.

For behavioral assay calibration, a fine thermocouple (70 µm diameter, type T, California Fine Wire Co, USA) was placed near the probe tip, at the approximate position of the larva during behavioral tests, to measure the actual temperature delivered to the surrounding air of larvae in experiments. If the measured temperature differed from the target temperature, the set temperature was adjusted accordingly. This setup ensured that the effective temperatures delivered to larvae matched the target temperatures.

#### Pad device

We developed a custom thermoelectric device to enable rapid and precise temperature control compatible with microscopy. The temperature-controlled stage consisted of a transparent diamond window (20 mm diameter, 250 µm thickness, k = 2000 W/m·K; Diamond Materials, Germany). The window was positioned between two cross-shaped copper parts with central openings. To minimize thermal contact resistance, indium sheets (k = 86 W/m·K) were inserted at the diamond–copper interface. Eight thermoelectric modules (1MDL06, TEC Microsystems, Germany) were integrated into the assembly, and a thermistor (GA10K3MCD1, TE Connectivity, USA), embedded in the copper part, provided feedback control via a TEC controller (TEC-1123, Meerstetter Engineering, Switzerland) powered by a 25.3 V DC supply. Heat generated at the hot side of the thermoelectric modules was dissipated through patterned copper heat-exchange blocks, which increased the effective surface area for thermal transfer. A temperature-controlled circulator chiller (Fisher Scientific Isotemp 6200 R28, USA) maintained the coolant of deionized water at 10 °C to ensure stable heat exchange during operation. The device provided stable control across 0–50 °C with a steady-state precision of ±0.1 °C.

For calcium imaging experiments, the pad device was calibrated using TEC Service software (version 3.00) to set the desired temperature. A Sylgard pad was attached to the surface of the temperature-controlled stage, and a fine thermocouple (70 µm diameter, type T, California Fine Wire Co, USA) was positioned at the approximate location of the larval head preparation during imaging to measure the actual temperature delivered to the larval tissue. This procedure ensured that the measured temperature at the sample matched the target setting.

### Behavioral Tests

Foraging third-instar larvae were used. All experiments were conducted on age- and size-matched larvae.

#### Cooling Assays

Cold/Cool behavioral assays were performed using a custom-made thermoelectric probe. The probe tip was pre-cooled to different temperatures and placed ∼100 μm in front of the larva’s head, adjacent to abdominal segment 4-7, or behind the tail (without touching the animal) to cool the surrounding air for 5 s during larval locomotion. An avoidance response was counted if the animal exhibited an aversive response specific to the target body region within the 5 s time window (Figures 1B and S2B). Behavioral analysis was performed double-blinded. Larvae’s head avoidance responses were recorded by a digital camera (EOS Rebel T5i, Canon, USA) inserted into the eyepiece of a stereo microscope (Olympus SZ61, USA). 6-10 larvae were tested on a single ∅ 35 mm grape-agar plate, and each animal was tested only once for behavioral responses specific to one body region. The response rate of all tested animals per genotype was calculated as a percentage, and chi-squared tests were performed for statistical analysis.

#### Heat Assays

Heat behavioral assays (Figures S5B and S5D) were performed using the same customized thermoelectric probe and recording settings as previously described for the cooling assays. The probe tip was pre-heated to 38°C and placed ∼100 μm in front of the larva’s head (without touching) to heat the surrounding air for 5 s. An avoidance response was counted if the animal exhibited head avoidance within the 5 s time window. The response rate of all tested animals per genotype was calculated as a percentage, and chi-squared tests were performed for statistical analysis.

#### Touch Assays

Touch behavioral assays (Figures S5B and S5D) were performed using a custom-made toothpick attached to a soft fiber and recording settings as previously described for the cooling and heat assays. The tip of the fiber was placed in light contact with the tip of the larva’s head for 2 s during larval locomotion. An avoidance response was counted if the animal exhibited head avoidance within the 5 s time window. The response rate of all tested animals per genotype was calculated as a percentage, and chi-squared tests were performed for statistical analysis.

#### Optogenetics

For optogenetic inhibition tests, *Drosophila* larvae were grown on food containing 1 mM all-trans-retinal (ATR) (A.G. Scientific). To inhibit candidate neurons, larvae carrying the UAS-GtACR1 transgene expressed GtACR1^18^ in C3da under 83B04-GAL4;^31^ in Cho under *iav*-GAL4; in C4da under *ppk*-GAL4;^30^ and in DOCCs under *R11F02*-GAL4.^10^ Larvae carrying only the UAS-GtACR1 transgene with no GAL4 driver (crossed to w^1118^) served as the negative control. During the test, 6-10 mid-third-instar larvae were placed on a ∅ 35 mm grape-agar plate. 25 μW/mm^2^ of 527-nm light was applied to activate GtACR1. Light intensity was measured with a power meter (PM16-122, Thorlabs, USA). The thermoelectric probe was set to different target temperatures (Figure 2A) and placed in front of the animal’s head for 5 s. Larvae’s head avoidance responses were recorded and scored as described for the cooling-evoked probe assay.

For optogenetic activation of C3da neurons, larvae were grown under the same ATR food conditions as described above for the optogenetic inhibition tests. Larvae grown on food without ATR served as the negative control. Larvae carrying the UAS-CsChrimson transgene expressed CsChrimson in C3da under 83B04-GAL4. During the test, a single mid-third-instar larva was placed on a ∅ 35 mm grape-agar plate and allowed to freely locomote. The plate was illuminated with a small-diameter (∼∅ 1 mm) circular spot of 655-nm red light (5 μW/mm^2^) emitted from a fluorescent microscope (SteREO Discovery.V8, Zeiss, Germany). To selectively activate C3da neurons in different regions of the larval body, the plate was moved into positions where the light selectively shone on the head, abdominal mid-section, or tail. To globally activate C3da neurons throughout the whole larval body, the diameter of the light was widened to ∼∅ 3 mm, and the plate was moved into positions where the light illuminated the entire larval body. Larvae’s behavioral responses were recorded via a mobile phone camera (iPhone 16, Apple, USA) and manually scored.

### Immunohistochemistry and Confocal Microscopy

Immunostaining of the head of third-instar larvae was modified from a previously reported method.^32,33^ The tip of 5-6 larvae were glued to the surface of a microscope glass slide using a mouth pipette in 1x phosphate-buffered saline (PBS), and a cut was made at the mouth hooks. After fixation in 4% formaldehyde for 30-40 min, larvae samples were washed with wash buffer for 10 times (2 min each). Samples were blocked in 100 μl of blocking buffer at room temperature for 1h before 1 μl of primary antibody was added. Samples were incubated with primary antibody for 4-6 hours. After incubation, samples were washed again, blocked in 100 μl of blocking buffer, before 1 μl of secondary antibody was added. Samples were incubated with secondary antibody overnight (12 hours) at 4°C. After incubation, samples were washed and mounted with DPX mounting media (Electron Microscopy Sciences, Hatfield, USA) before imaging.

Primary antibodies used include chicken anti-GFP (1:2500, Aves Laboratories, RRID: AB_2307313). Secondary antibodies used include anti-chicken Alexa Fluor 488 (1:500, Jackson ImmunoResearch, RRID: AB_2340375). Confocal imaging was performed on a Leica SP8 confocal system equipped with a resonant scanner using the 20x lens. Images were collected with z stacks of 1.0-μm-step size. The acquired three-dimensional images were projected into two-dimensional images using the maximum projection method.

### In Vivo Calcium Imaging

Larvae anesthetized by chloroform were placed in a droplet of HL3 imaging buffer on a sylgard-coated cover glass. The head was cut below the VNC (ventral nerve cord) using dissection scissors, which left only the aC3da neurons intact for imaging, and subsequently glued to the sylgard surface using a mouth pipette (Figure S3A). Head dissociation from the body was necessary for secure gluing. The remainder of the larval body was discarded, and the solution was replaced with fresh buffer consisting of modified hemolymph-like 3 (HL3) saline (70 mM NaCl, 5 mM KCl, 0.5 mM CaCl_2_, 20 mM MgCl_2_, 5 mM trehalose, 115 mM sucrose, 5 mM HEPES, and 7 mM glutamate, pH 7.2).^34,35^ The sylgard pad with the dissociated larval head was adhered to the cooling chamber of a custom-built thermoelectric pad device, where different set temperatures were reached. Cool and cold-evoked calcium responses were recorded from a stereo microscope (Stemi SV11 Apo Stereo Motorized Microscope, Zeiss, Germany) using a 20x/NA0.60 DRY objective lens.

The thermoelectric pad has a temperature ramp rate of 5°C/s. The temperature was initially set at 23°C; after the calcium trace stabilized, the temperature was cooled to 18°C over 30 s, heated back to 23°C over 15 s, cooled again to 13°C over 30 s, then heated back to 23°C (Figure S3B). In an adjusted temperature paradigm, the temperature was initially set at 23°C; after the calcium trace stabilized, the temperature was cooled to 18°C over 30 s, and heated back to 23°C (Figure S3C).

For both temperature protocols, we scored the percentage change in the maximum intensity of GCaMP fluorescence.

### In Vitro Calcium Imaging (cell culture)

Calcium imaging of CHO cells was performed as previously described.^23^ Cells were maintained in DMEM/F12 supplemented with 10% fetal bovine serum at 37°C in a 5% CO₂ incubator. For imaging, cells were plated on poly-D-lysine–coated coverslips in 35 mm dishes and transfected for 4 hours using Lipofectamine 3000 (ThermoFisher), with a lipid-to-DNA ratio of 3:1 and a total of 1.0 μg DNA per dish. After 18 hours, cells were incubated with 2.5 μM Fura-2 AM and 0.2% Pluronic F127 in HBSS for 30 minutes at room temperature. Cells were then transferred to imaging solution (10LmM HEPES, pH 7.4, 145 mM NaCl, 5 mM KCl, 1.2 mM MgCl₂, 2.5 mM CaCl₂, and 10 mM glucose) and incubated for an additional 30 minutes before recording.

Imaging was performed on an Olympus IX73 inverted microscope with a 20x objective. Fluorescence signals were acquired using a Hamamatsu ORCA-Flash 4.0 sCMOS camera and MetaFluor software (Molecular Devices). The imaging solution was perfused to the cells, and the solution’s temperature was regulated using a Bipolar In-line Cooler/Heater (SC-20, Warner Instruments). Cells were initially exposed to 25°C, and after stabilization over 30-50 s, the temperature was cooled to 18 or 13°C over 200 s, then returned to 25°C (Figure S7). We scored the percentage change in the peak Fura-2 fluorescence ratio (ΔR/R_0_), where R represents the 340/380 nm excitation ratio.

### Statistical Analysis

All statistical analyses were performed using GraphPad Prism 9.0 software. Statistical methods, error bars, and p-values are indicated in the figure legends. Sample numbers are indicated in the figure legends. Chi-squared tests, Unpaired Student’s t-tests, Paired Student’s t-tests, and One-Way ANOVA with various methods of multiple comparisons post hoc analysis were applied separately to analyze data, and further described in the figure legends. For all statistical analyses: ns for p>0.05, * for p<0.05, ** for p<0.01, *** for p<0.001, **** for p<0.0001.

## Supporting information

Video S1. Head avoidance of cold

Video S2. Tail unresponsiveness to cold

Video S3. Mid-section avoidance of cold

Video S4. Activation of anterior C3da neurons

Video S5. Activation of mid-section C3da neurons

Video S6. Activation of posterior C3da neurons

Video S7. Full-body activation of C3da neurons

## ACKNOWLEDGEMENTS

We thank Shuhao Wan, Ty Hergenreder, and Can Wang for technical support. This work was supported by grants from the National Institutes of Health (NIH) to B.Y. (R01NS128500) and the National Research Foundation of Korea (NRF) grant funded by the Korean government (MSIT) to G.-H.K. (2021R1A2C2014512). *Drosophila* Stocks from the Bloomington *Drosophila* Stock Center (NIH P40OD018537) were used in this study.

## AUTHOR CONTRIBUTION

L.A.X., E.A.R., G.-H.K., and B.Y. conceived the project. L.A.X. performed the *in vivo* experiments and analyzed the data with guidance and assistance from R.L. and E.A.R. X.B. performed the *in vitro* cell culture experiments and analyzed the data. H.-J.S. and G.-H.K. contributed to the design and fabrication of the cooling devices. L.A.X. and B.Y. wrote the manuscript with input from all other authors.

