## Supplemental Figures for "Cold sensing by a glutamate receptor drives avoidance behavior in *Drosophila* larvae"

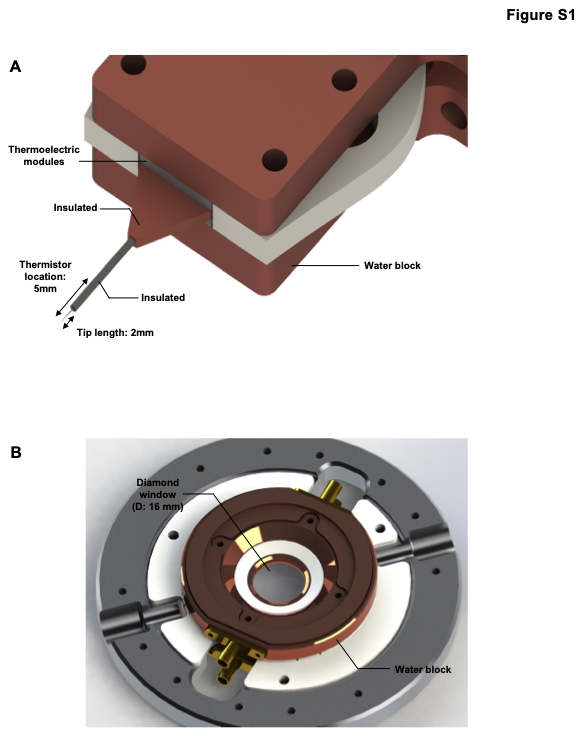

**Figure S1. Schematic illustrations of the thermoelectric cooling devices**

(A) Probe device for localized temperature control. The nickel-plated tungsten probe tip was insulated along most of its length, with 2 mm exposed at the tip for cooling. A thermistor was positioned 5 mm from the tip to provide temperature feedback. The copper component was insulated to enhance cooling performance and stability.
(B) Pad device for calcium imaging experiments. The temperature-controlled stage consisted of a 20 mm diameter diamond window, thermally regulated via heat exchange at its 2 mm edges, enabling central temperature control with a precision of ±0.1°C.

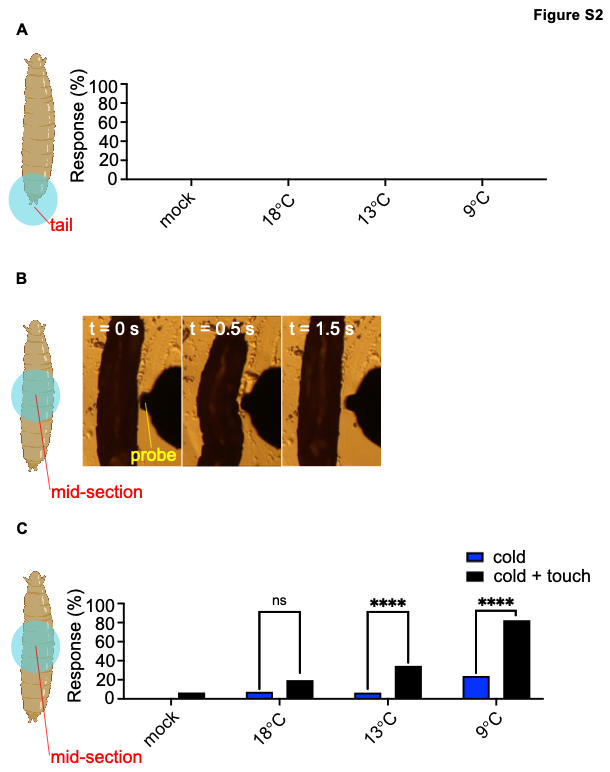

**Figure S2. The mid-section and tail of *Drosophila* larvae are less sensitive to cold**

(A) An absence of observable cold-evoked responses in the tail. Wild-type Canton-S larvae were tested at mock (ambient) temperature (n = 51), 18°C (n = 63), 13°C (n = 61), and 9°C (n = 40).
(B) Representative images of the mid-section avoidance behavioral sequence.
(C) Simultaneous cold and mechanical stimuli induce more robust mid-section avoidance responses than those induced by cold stimuli alone. Wild-type Canton-S larvae were tested with a non-contacting (cold alone) and a contacting probe (cold and touch). Non-contacting probe: mock (n = 41), 18°C (n = 40), 13°C (n = 76), and 9°C (n = 50). Contacting probe: mock (n = 75), 18°C (n = 61), 13°C (n = 49), and 9°C (n = 40). Chi-squared test. ****p<0.0001.

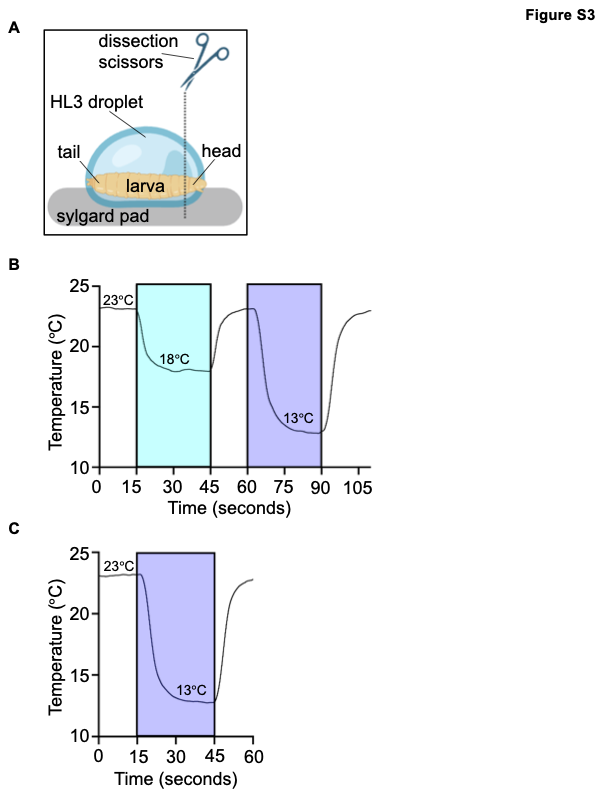

**Figure S3. Methods for calcium imaging of aC3da neurons**

(A) Schematic of larval head prep for calcium imaging.
(B) A thermoelectric pad device with a ramp rate of 5°C/s was used to provide thermal stimulation during calcium imaging of aC3da neurons in larval heads. For double-stimulation paradigms (Figure 4), an initial temperature of 23°C was set, and neuronal responses were allowed to stabilize for 15 s. The temperature was cooled to 18 °C over 30 s, heated back to 23°C over 15 s, cooled again to 13°C over 30 s, then heated back to 23°C.
(C) For single-stimulation paradigms (Figure S5), an initial temperature of 23°C was set, and neuronal responses were allowed to stabilize for 15 s. The temperature was cooled to 13°C over 30 s, then heated back to 23°C.

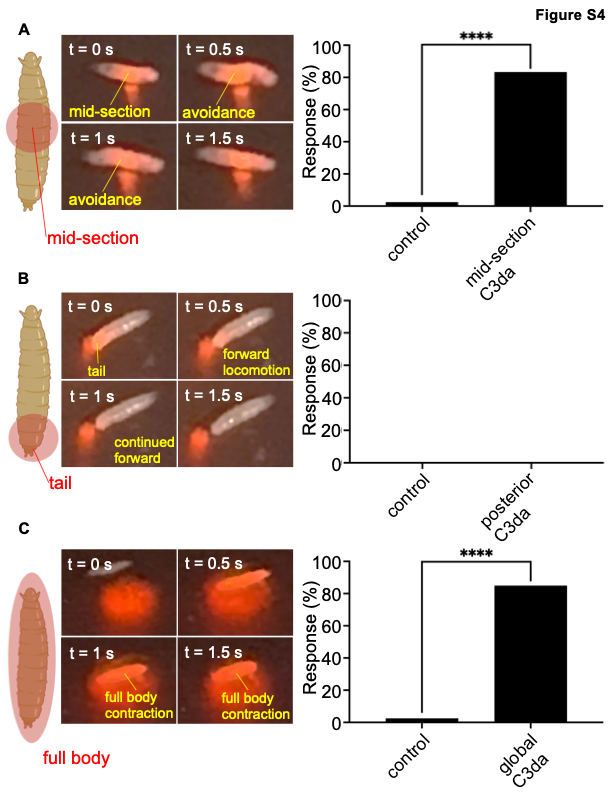

**Figure S4. Optogenetic activation of 83B04-positive (C3da) neurons in different body regions of *Drosophila* larvae elicits distinct behavioral responses**

(A) Selective optogenetic activation of 83B04-GAL4 labeled neurons in the larval abdominal mid-section elicits mid-section avoidance behavior during forward peristalsis, where the abdominal midsegments bend away from the light in an act of evasion. Larvae grown on non-ATR-containing food subjected to the same illumination conditions served as the negative control. Chi-squared test. Bar graph, n = 41 (control), n = 42 (mid-section C3da). ****p<0.0001.
(B) Selective optogenetic activation of 83B04-GAL4 labeled neurons in the larval tail elicits no noticeable behavioral responses during forward peristalsis. Larvae grown on non-ATR-containing food subjected to the same illumination conditions served as the negative control. Chi-squared test. Bar graph, n = 44 (control), n = 43 (posterior C3da).
(C) Optogenetic activation of 83B04-GAL4 labeled neurons throughout the entire larval body elicits full body contraction behavior, where the head and tail withdraw simultaneously. Larvae grown on non-ATR-containing food subjected to the same illumination conditions served as the negative control. Chi-squared test. Bar graph, n = 40 (control), n = 40 (global C3da). ****p<0.0001.

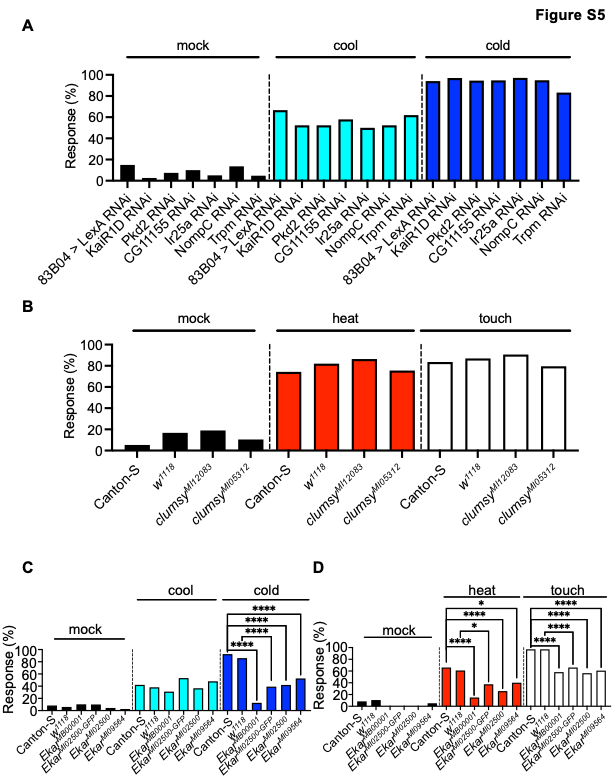

**Figure S5. Head avoidance responses of candidate genetic mutants and RNAi knockdowns to thermal and/or mechanical stimuli**

(A) RNAi-mediated knockdown of candidate receptor-encoding genes in 83B04-positive C3da neurons does not affect head responses to cool and cold temperatures. Chi-squared test. Bar graphs, n $\geq$40 per genotype.
(B) The head of *Clumsy* mutants responds normally to heat and mechanical touch. Chi-squared test. Bar graphs, n $\geq$40 per genotype.
(C) Cool and cold-elicited head avoidance responses of *Ekar* mutants. Each *Ekar* mutant was compared to its corresponding genotypic background (either Canton-S or *w^1118^*). Chi-squared test. Bar graph, n $\geq$ 40 per genotype. ****p<0.0001.
(D) Heat (34 °C) and touch-elicited head avoidance responses of *Ekar* mutants. Each *Ekar* mutant was compared to its corresponding genotypic background (either Canton-S or *w^1118^*). Chi-squared test. Bar graph, n $\geq$ 40 per genotype. *p<0.05 and ****p<0.0001.

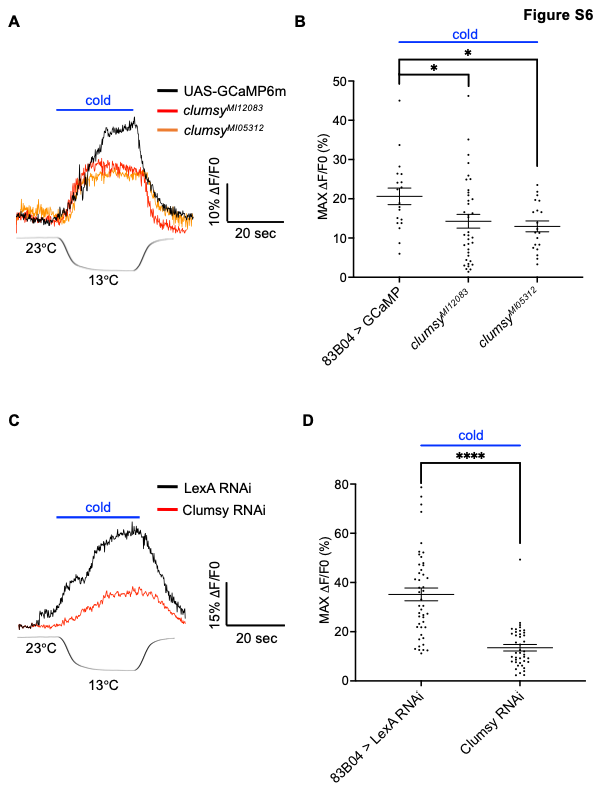

**Figure S6. Cold sensitivity of aC3da neurons requires Clumsy**

(A) Representative traces of cold-induced calcium transients of aC3da neurons in GCaMP-expressing controls and *Clumsy* mutants. Length of the temperature label above the trace corresponds to the duration of cold stimulation.
(B) Maximum $\Delta$F/F_0_ of individual aC3da neuronal clusters is plotted on the graph and used for statistical analyses (each dot indicates one cluster). Control (n = 19), *Clumsy^MI12083^* (n = 37), *Clumsy^MI05312^* (n = 20). One-Way ANOVA with Tukey multiple comparisons post hoc analysis. Error bars represent SEM. *p<0.05.
(C) Representative traces of cold-induced calcium transients of aC3da neurons in GCaMP-expressing controls and with *Clumsy* RNAi-mediated knockdown. Length of the temperature label above the trace corresponds to the duration of cold stimulation.
(D) Maximum $\Delta$F/F_0_ of individual aC3da neuronal clusters is plotted on the graph and used for statistical analyses (each dot indicates one cluster). Control (n = 46) and clumsy RNAi (n = 42). Unpaired Student’s t-test. Error bars represent SEM. ****p<0.0001.

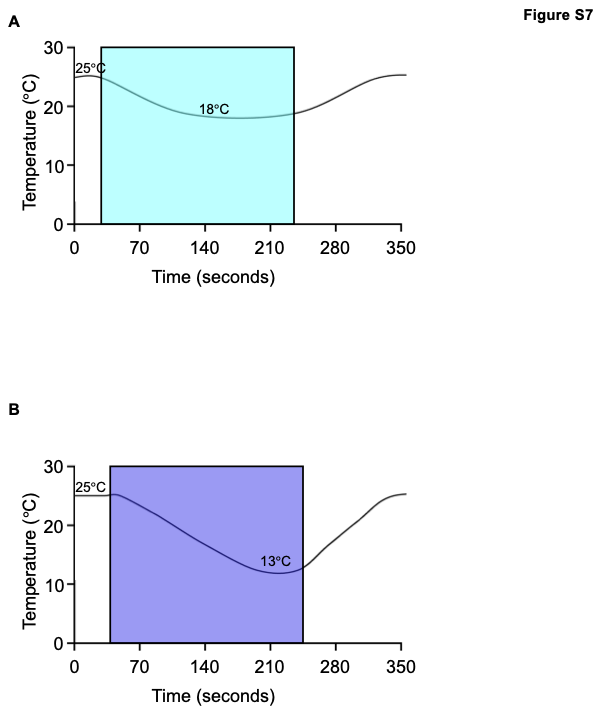

**Figure S7. Temperature protocol for Fura-2 imaging of CHO cells**

(A) A perfusion system regulated via a Bipolar In-line Cooler/Heater with a ramp rate of 0.08°C/s was used to provide thermal stimulation during calcium imaging of transfected CHO cells. For examining cool-evoked calcium transients, an initial temperature of 25°C was set, and responses were allowed to stabilize for 30-50 s. The temperature was cooled to 18°C over ~200 s, then heated back to 25°C.
(B) A perfusion system regulated via a Bipolar In-line Cooler/Heater with a ramp rate of 0.08°C/s was used to provide thermal stimulation during calcium imaging of transfected CHO cells. For examining cold-evoked calcium transients, an initial temperature of 25°C was set, and responses were allowed to stabilize for 30-50 s. The temperature was cooled to 13°C over ~200 s, then heated back to 25°C.
